# Brain geometry matters in Alzheimer’s disease progression: a simulation study

**DOI:** 10.1101/2020.07.24.220228

**Authors:** Masoud Hoore, Jeffrey Kelling, Mahsa Sayadmanesh, Tanmay Mitra, Marta Schips, Michael Meyer-Hermann

## Abstract

The Amyloid cascade hypothesis (ACH) for Alzheimer’s disease (AD) is modeled over the whole brain tissue with a set of partial differential equations. Our results show that the amyloid plaque formation is critically dependent on the secretion rate of amyloid β (Aβ), which is proportional to the product of neural density and neural activity. Neural atrophy is similarly related to the secretion rate of Aβ. Due to a heterogeneous distribution of neural density and brain activity throughout the brain, amyloid plaque formation and neural death occurs heterogeneously in the brain. The geometry of the brain and microglia migration in the parenchyma bring more complexity into the system and result in a diverse amyloidosis and dementia pattern of different brain regions. Although the pattern of amyloidosis in the brain cortex from in-silico results is similar to experimental autopsy findings, they mismatch at the central regions of the brain, suggesting that ACH is not able to explain the whole course of AD without considering other factors, such as tau-protein aggregation or neuroinflammation.

## Introduction

Alzheimer’s disease (AD) is the most common dementia [4] from which around one third of people older than 85 years suffer [25]. This number is increasingly alarming as life expectancy grows, while a treatment has not been found yet [4]. The Amyloid cascade hypothesis (ACH) [24] is the strongest among its peers explaining the course of the disease through a cascade of events [35].

Amyloid-*β* (A*β*) is the leading agent in AD according to the ACH [24, 32, 60], causing the other events to follow after. A*β* is secreted into the interstitial fluid (ISF) from the neural synapses by abnormal cleavage of amyloid precursor protein (APP) [64, 12, 62, 57]. APP can be cleaved by *γ*-secretase from one side, and either *α*- or *β*-secretase on the other. If cleaved by *α*-secretase, APPs*α* is produced which is known to be neurotrophic and neuroprotective [51]. However, when cleaved by *β*-secretase, A*β* is excreted. The activity of *β*-secretase enzymes, BACE1, increases with age [17, 74, 18, 43], implying that more A*β* is secreted from the neural synapses. The concentration of A*β* has been shown to be correlated with the age as well [55, 19]. More A*β* production due to higher activity of BACE1 means in addition that fewer APPs*α* are produced, hindering their neurotrophic and neuroprotective effect [39]. On the other hand, more production of A*β* in the interstitial fluid (ISF) increases the chance that they may aggregate in plaques.

The brain immune system, namely microglia and astrocytes, together with the cerebrospinal fluid (CSF) try to clear the produced A*β*. Microglia not only clear the brain from soluble A*β* through micropinocytosis [45, 71, 40], but also migrate toward the amyloid plaques for digesting them through phagocytosis [10, 8]. Similarly, astrocytes migrate toward the plaques [38] for separating the inflamed zone from the healthy tissue [65]. Soluble A*β* can also discharge mechanically from the cerebrospinal fluid (CSF)[6, 46, 63] or meningeal lymphatics [14, 30]. Nevertheless, the clearance of A*β* cannot tolerate production rates beyond a critical value [29]. This leads to the emergence of a critical production rate (A*β* concentration) over which the plaques form [29, 19].

The AD process from A*β* accumulation to the onset and symptomatic dementia stages is a complicated scenery of the interplay among the brain immune system, neural activity, and aggregation of A*β* into plaques. In addition to such complexity, the course of the disease spans over years to decades from the onset till the symptomatic stage. Consequently, it is tedious to study the disease systematically and check whether a proposed treatment is working or not. Modeling may come very useful, however, for understanding the disease and implementing the suggested interventions and verify their efficacy. We propose a tempospatial mathematical approach with partial-differential equations (PDEs) for this process and investigate the effect of neural density/activity and the brain tissue structure on AD by computer simulations.

Several models for AD have been proposed and studied so far. On a small scale, amyloid plaque formation and its interaction with microglia has been modeled using agent-based simulations [15]. Larger mean-field models have also been proposed for AD, using partial and ordinary differential equations (PDEs and ODEs) [23, 58]. Recently, more sophisticated models have tried to bring the brain structure into play. Bertsch et al. [7] in 2016 have introduced a system of PDEs to model the onset and progression of AD. Although they use a 2D space with simple boundary conditions, their model is able to qualitatively capture the principal behavior of the brain in AD such as the ACH. Other models argue that the diffusion of prions is mainly in the direction of neural fibers and find matching atrophy patterns between the model and real AD patients [59, 72]. In our model, we approximate the neural density from structural MR images, and presume that A*β* secretion is proportional to it. Considering the real brain tissue structure from MRI data as the boundary conditions for our system of PDEs, our simulation results show, that not only the neural density, but also the brain geometry and tissue structure is important in determining where amyloid plaques form. The model shows how the interplay between the neurons, prions, the brain structure and its immune system affects the progression of AD. Based on our model, we argue that AD is a result of a phase change in A*β* physical form from soluble to the plaques, driven by its production from neural synapses. In consequence, neural atrophy is related to many factors and occurs heterogeneously in the brain. We finally provide which brain lobes are under higher probability of plaque formation and subsequent atrophy. According to our results, we argue that ACH is not sufficient to predict the atrophy in the brain if only amyloids are considered as the main culprits of the disease.

## Model

Our equation-based model is described by Eqs. 1 to 7 as depicted in Fig. 1.

**Figure 1:**
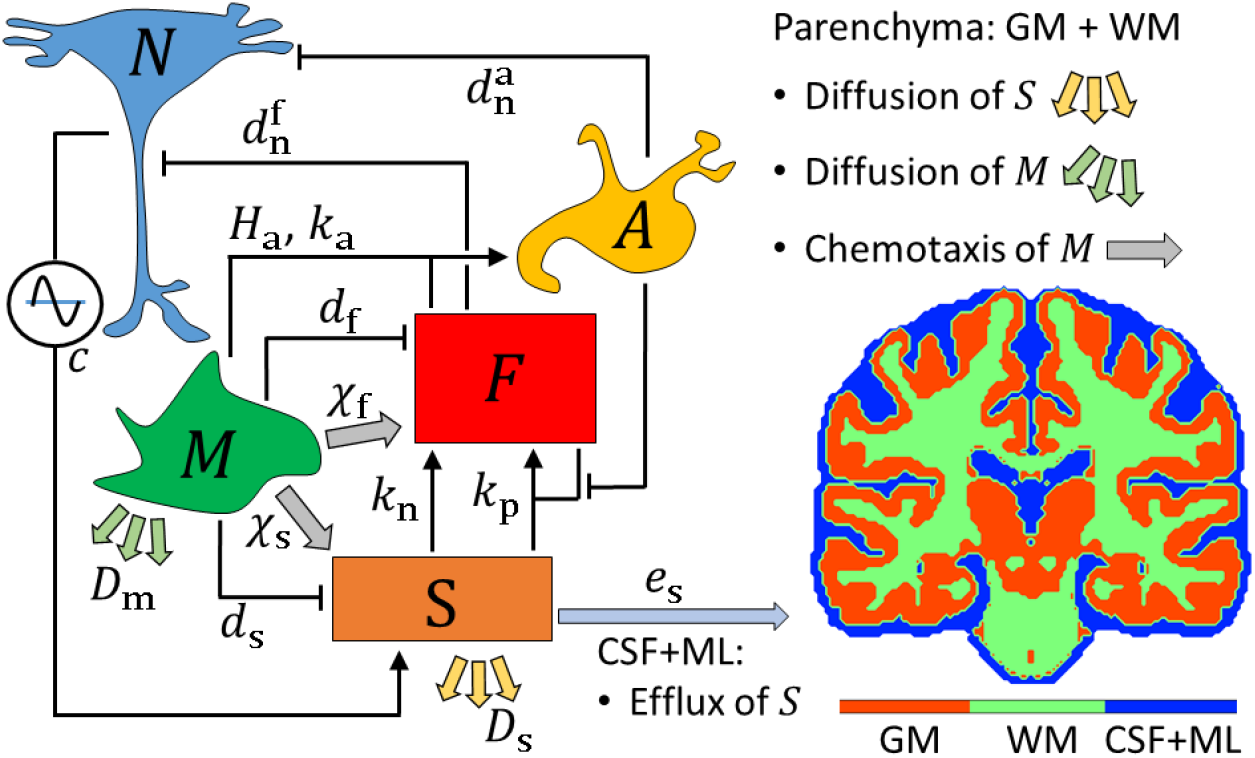
The network of interactions for the AD model. Microglia (*M*) randomly wander in the parenchyma, do chemotaxis toward soluble (*S*) and fibrillized A*β* (*F*), to clear them. *S* is produced by neurons (*N*), due to neural activity *c*. It diffuses in the parenchyma and discharges through the cerebrospinal fluid (CSF) or meningeal lymphatics (ML), surrounding the parenchyma. *F* forms out of *S* through nucleation and polymerization processes. Astrocytes get activated by the signaling from microglia when they perform phagocytosis on the amyloid plaques. When astrocytes get highly activated, they form scars around the plaques. The activation of astrocytes is known as astrogliosis *A*. It is assumed that both *F* and *A* signal the death of neurons. Brain parenchyma comprises the white matter (WM) and the grey matter (GM), as shown in a coronal cross section. *M*, *F*, and *A* are zero in CSF. The brain tissue is taken from MR images of Ref. [16] and the neural density is estimated from the MRI with respect to Ref. [5].

### Production, fibrillization, and clearance of A*β*

Soluble A*β* is secreted from the synapses due to synaptic activity [12, 64, 39]. It may form fibrillar amyloid plaques or get discharged from the brain through the cerebrospinal fluid (CSF) [6, 46, 63] or meningeal lymphatics [14, 30]. The changes of the concentration of soluble and fibrillar A*β* are governed by

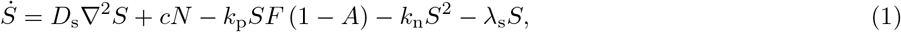

and

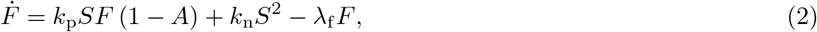

where *S* and *F* stand for the concentration of A*β* oligomers in soluble ([sA*β*]) and fibrillar plaque ([fA*β*]) form, respectively. The term *D*_s_▽^2^*S* in Eq. 1 represents the diffusion of sA*β* in the brain parenchyma with diffusivity *D*_s_, the term *cN* is the production of sA*β* from neurons. Here, *N* stands for the neural density, and *c* is a measure of neural activity. An accurate distribution of *N* and *c* in space and time is not available so far. We assume that the distribution of *N* in space is proportional to the structural MR image intensities. The mean neural density in brain parenchyma is estimated as 〈*N*〉 = 1.3^7^ mL^*−*1^ according to Ref. [5]. *c* seems to be the most complicated variable in the model as it is a function of both space and time due to different activities at different brain lobes and different times in the circadian rhythm. *c* is assumed to be constant and homogeneous throughout the brain. We estimate it to be 〈*c*〉 = 2.0 10^*−*11^ *μ*M mL day^*−*1^ from our most realistic simulation results. It is noted however that *c* can vary for different tissues of the brain, e.g. Moghekar and colleauges in 2011 have estimated the amount of A*β* secretion from each neuron in vitro [52], arriving at *c* ≈ 4.3 × 10^*−*10^ *μ*M · mL · day^*−*1^.

The terms *k*_p_*SF* (1 − *A*) and *k*_n_*S*^2^ represent the formation of fibrillar plaques through two distinct mechanisms, respectively polymerization and nucleation. In the absence of amyloid plaques, soluble A*β* oligomers form small nuclei with a tiny nucleation rate *k*_n_. The growth of the nuclei occurs much faster with a polymerization rate *k*_p_ which is greater than *k*_n_ [44, 29]. The growth of amyloid plaques can be prohibited by astrocytes through astrogliosis and scar formation, represented by *A* (see Eq. 6).

The term *λ*_s_*S* signifies the clearance of sA*β* by either microglia through macropinocytosis with the rate *d*_s_ [45, 71, 40], or sA*β* efflux through the CSF or meningeal lymphatics (ML) with rate *e*_s_ [6, 46, 14, 30]. In other words,

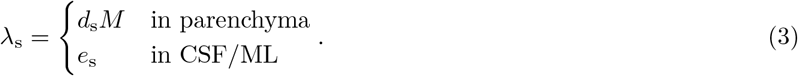

*M* represents the number density of the microglial cells. Similarly, *λ*_f_ in Eq. 2 is

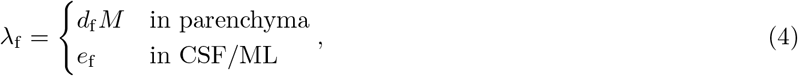

where *d*_f_ signifies the phagocytic rate of fA*β* by microglia. Fibrillization of A*β* in CSF is ignored in the model, i.e. *F* = 0 in CSF. A recent investigation on a simplified version of Eq. 1 and 2 shows that the fibrillization of sA*β* to fA*β* is mainly triggered by the production rate *cN*. We also found that there is a critical production rate beyond which the fibrillization occurs. The other parameters of the system regulate this critical rate [29].

### Microglia movement and chemotaxis

It is known that microglia actively move in the brain parenchyma (white (WM) and grey matter (GM)) using their processes [56], and concentrate on the regions with high amyloid plaque burden [1, 42, 71, 31]. The change of microglial cell density in time is given by

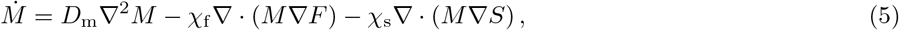

in which *D*_m_, *χ*_f_, and *χ*_s_ stand for the effective diffusivity of the microglia due to their random-walk movement, and chemotactic sensitivity [37] toward *S* and *F*, respectively. It is evident from the experiments that microglia gather at the spots of amyloid plaques [71, 31]. We do not know whether such migration is a chemotactic movement toward the gradient of either sA*β*, fA*β*, or both. Therefore, a chemotactic movement toward both forms of A*β* is modeled. The microglial proliferation is neglected because it occurs on a shorter time scale with shorter settlement time. Eq. 5 is valid only in the brain parenchyma.

### Astrogliosis

The whole process of the astrocyte response to injury or disease in CNS is called reactive astrogliosis [65]. Astroglyiosis can be triggered by amyloid plaques [11, 22] and results in scar formation by the highly activated astrocytes at their boundaries [66, 48]. Through astrogliosis, astrocytes emit many signaling factors, triggering microglia and neurons. Scar formation occurs at the highest level of astrogliosis, which limits the growth of amyloid plaques by creating a physical obstacle between the plaques and their surrounding soluble A*β* oligomers. Although the scars protect neurons from getting intoxicated by A*β* plaques, they may be detrimental for neurons through other routes. All normal interactions between neurons and microglia are destroyed at the scar site [66], implying that dementia may result from a progressive scar formation.

We simplify such a complex interactive network by introducing a dimensionless number between zero and one, 0 ≤ *A* ≤ 1, as a representation of the extent of astrogliosis. *A* ≪ 1 means a healthy state of the brain, and *A* ∼ 1 implies a highly inflamed state with strong astrogliosis. The scar formation is represented by multiplying a factor (1 − *A*) to the polymerization term in Eq. 1 and 2.

Astrogliosis is assumed to be triggered by microglia when they perform phagocytosis on amyloid plaques and secret neuroinflammatory signals [50, 27]. Thus, astrogliosis is supposed to be related to both *F* and *M*. The process should saturate, because once all astrocytes are activated, they cannot get further activated. We consider a Michaelis-Menten kinetics [49, 33] for the activation of astrocytes, as

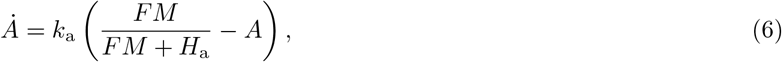

where *k*_a_ is the astrogliosis rate, and *H*_a_ is the Michaelis constant of this process.

### Neural death

According to the ACH, amyloid plaques either directly or indirectly cause the death of neurons [24, 32, 60]. Some studies suggest that the neurons are not directly intoxicated by amyloid plaques [69], but they are rather affected by the subsequent processes triggered by plaques, such as the formation of the neurofibrillary tangles inside neurons [20]. It is also suggested, that neuroinflammation may cause neural stress and contribute to their death. In consequence, we assume that neurons die due to both astrogliosis and fibrillization, as

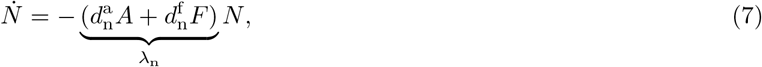

with the death/decay rates 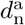 and 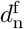.

### Parameter estimation and the simulation method

In order to understand the overall behavior of the system, the spatial derivatives are neglected from the equations. Such an approach helps us in finding the correct range of parameters for a realistic case. The results of this approach are only presented in the supplementary materials. Afterwards, the spatial derivatives and the brain tissue structure are included in the calculations and the model is solved for a more specific parameter space. More details on the simulation methods are given in the supplementary materials.

## Results & Discussion

### Brain geometry uncouples neural density and neural atrophy

Naively speaking, more neurons would result in more amyloid production, therefore more amyloidosis and more atrophy. However, since the brain has a broad distribution of neural density and activity, a wide range of *cN*_o_ (shown in Fig. S2) should occur in a single brain. That means, some parts of the brain may be under higher amyloidosis pressure, but the other parts be more relaxed (more on this will be explained in Fig. 7). A manifestation of this could be the fact that the most active regions of the brain, represented by the default mode network (DMN), contain the highest burden of amyloid plaques and are the first regions of dementia in AD [21, 26, 67, 53].

On the other hand, soluble amyloid oligomers can diffuse in the brain parenchyma. The estimations from the mean-field analysis (see supplementary materials) shows that A*β* is mainly cleared through the CSF rather than by microglia. However, sA*β* should have enough time to diffuse out to the CSF in order to be cleared effectively. That means the regions of the brain, which produce high amounts of amyloids and are far from the CSF, possess a high risk of fibrillization. Moreover, the migration of microglia toward the plaques, and their random-wandering in the brain add to the complexity of the system by making the system highly heterogeneous.

The z-score maps, the normalized spatial distribution, of all the system parameters provide a better view of what may actually happen in the brain (Fig. 2). The time evolution of the variable means (Fig. 3) is also a useful measure of how the disease progresses. The z-score of variable *x* is defined as

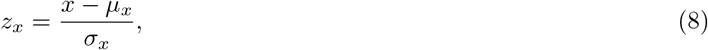

at each position, where *μ*_*x*_ and *σ*_*x*_ are the mean and standard deviation of variable *x* throughout the brain.

**Figure 2:**
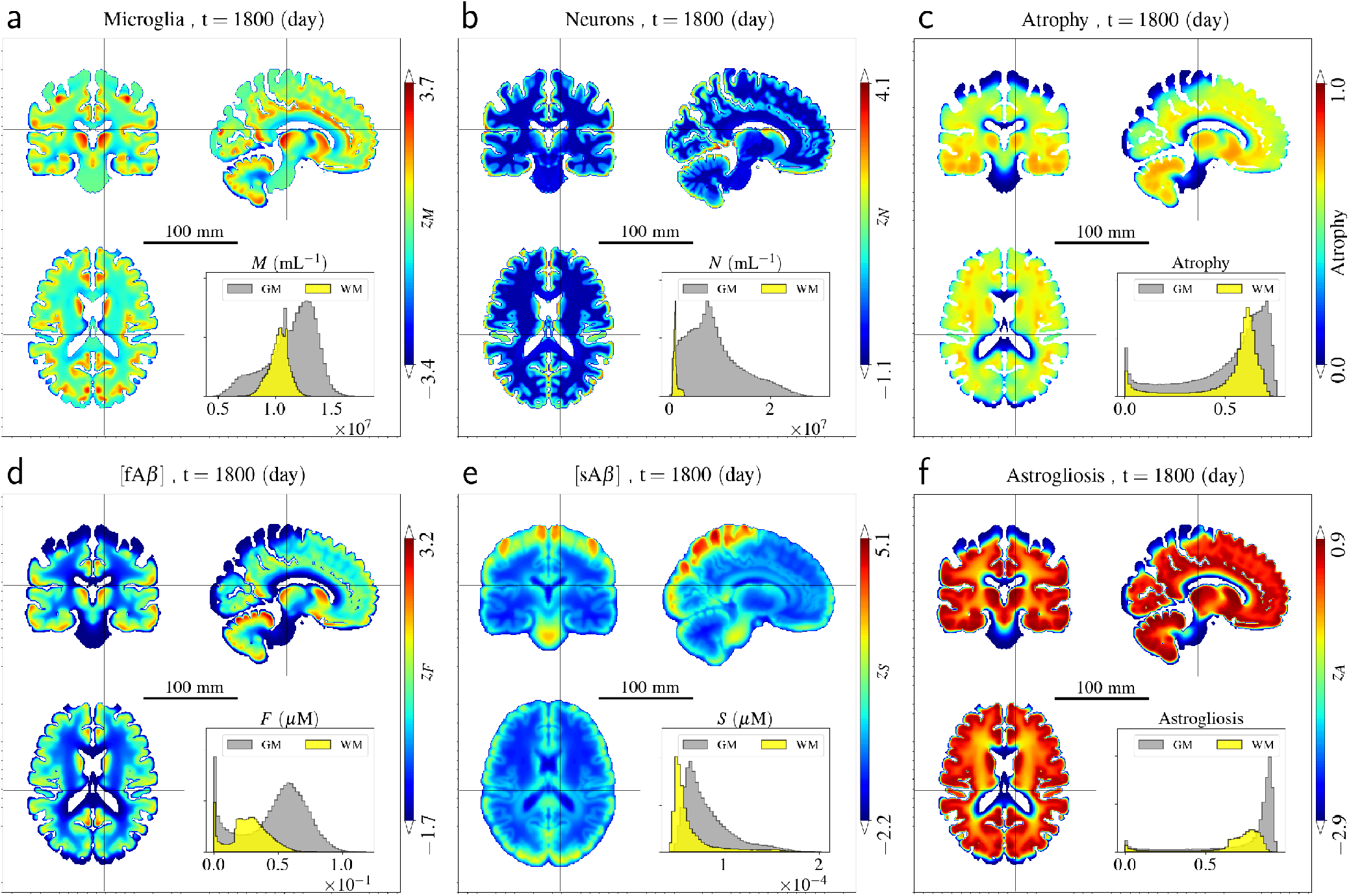
Spatial map of the system variables at a single time *t ≈* 5 yr. The panels show the z-score map of the system variables. The inset diagrams show the histogram of each paramer in WM and GM. For the full time evolution of the z-score maps, see SM1.

**Figure 3:**
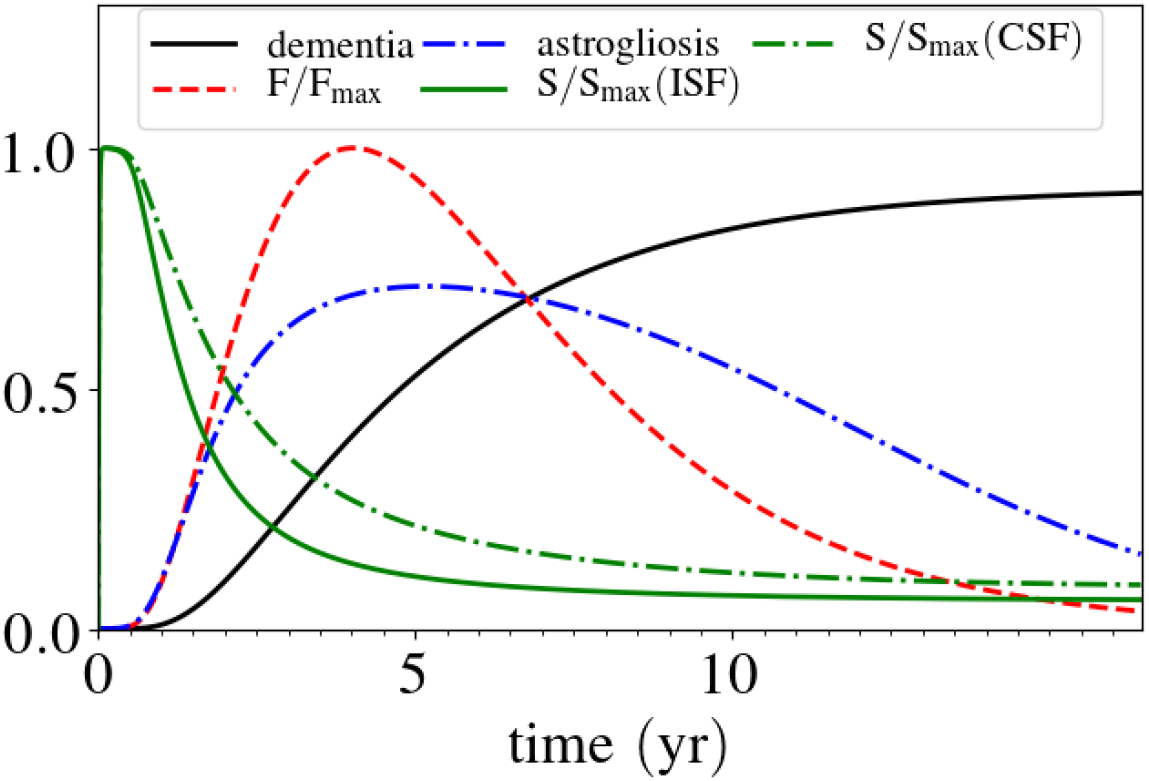
Timeline of the mean of the system variables, dementia (1 *− N/N*_o_), astrogliosis *A*, [sA*β*] *≡ S*, and [fA*β*] *≡ F*. The initial time is when the homeostatic state of the brain switches to fibrillization dominance.

It is noted that the initial time (*t* = 0) stands for the time after which the homeostatic state of the system switches from normal soluble amyloid dominance toward the amyloid fibrillization dominance. What the absolute value of this time is for each individual, meaning the time after birth, remains a big question. Definitely, it is around 70 to 80 years after birth on average. However, it is out of scope of this study.

Here, dementia is defined as the total loss of neurons - atrophy, normalized by the original number of neurons, *i.e.* dementia = *∫*atrophy · d*v* = *∫*(1 − *N*(*t*)*/N*(0))d*v*. This term should be distinguished from the conventional dementia score. The onset of AD can be defined as the time point after which the dementia surpasses a threshold. Such a threshold should be different for each individual due to their brain reserve capacity (BRC) [68]. The BRC could be associated with the total number of neurons or the resilience of the brain neural network against dementia both of which should be related to each other. Since the dementia is normalized by the initial (original) neural density (*N*_o_), one cannot assign the onset of AD to a unique threshold value. Still, however, Fig. 3 shows that before the onset of AD, sA*β* drops substantially from its saturated point. In healthy condition, sA*β* are mostly generated at the GM, and diffuse out to the CSF. However, they may sediment into amyloid plaques on their way if their production rate is higher than a critical value [29].

As *S* fades away, *F* rises up and triggers astrogliosis *A*. *A* soon takes the whole brain parenchyma, as shown in Fig. 2-f and SM1. Consequently, further fibrillization is blocked and *F* decays slowly by microglia. However, the clearance of *F* takes much longer time than their aggregation. Meanwhile, neurons die because of intoxication by *A* and *F*. The burden of amyloid plaques *F*, neural atrophy, microglial density *M*, and astrogliosis peak at the highest neural density/activity regions (compare Fig. 2-a, c, d and f with Fig. S6).

The time evolution of *F* demonstrates that the first amyloid plaques form in the central regions of the brain, followed by fibrillization in the grey matter cortex (see SM1). Fig. 4 also shows that the last region of the brain taken by amyloid plaques is the parietal lobe. These results are in agreement with the reported amyloid burden in the brain of AD patients [9, 34].

**Figure 4:**
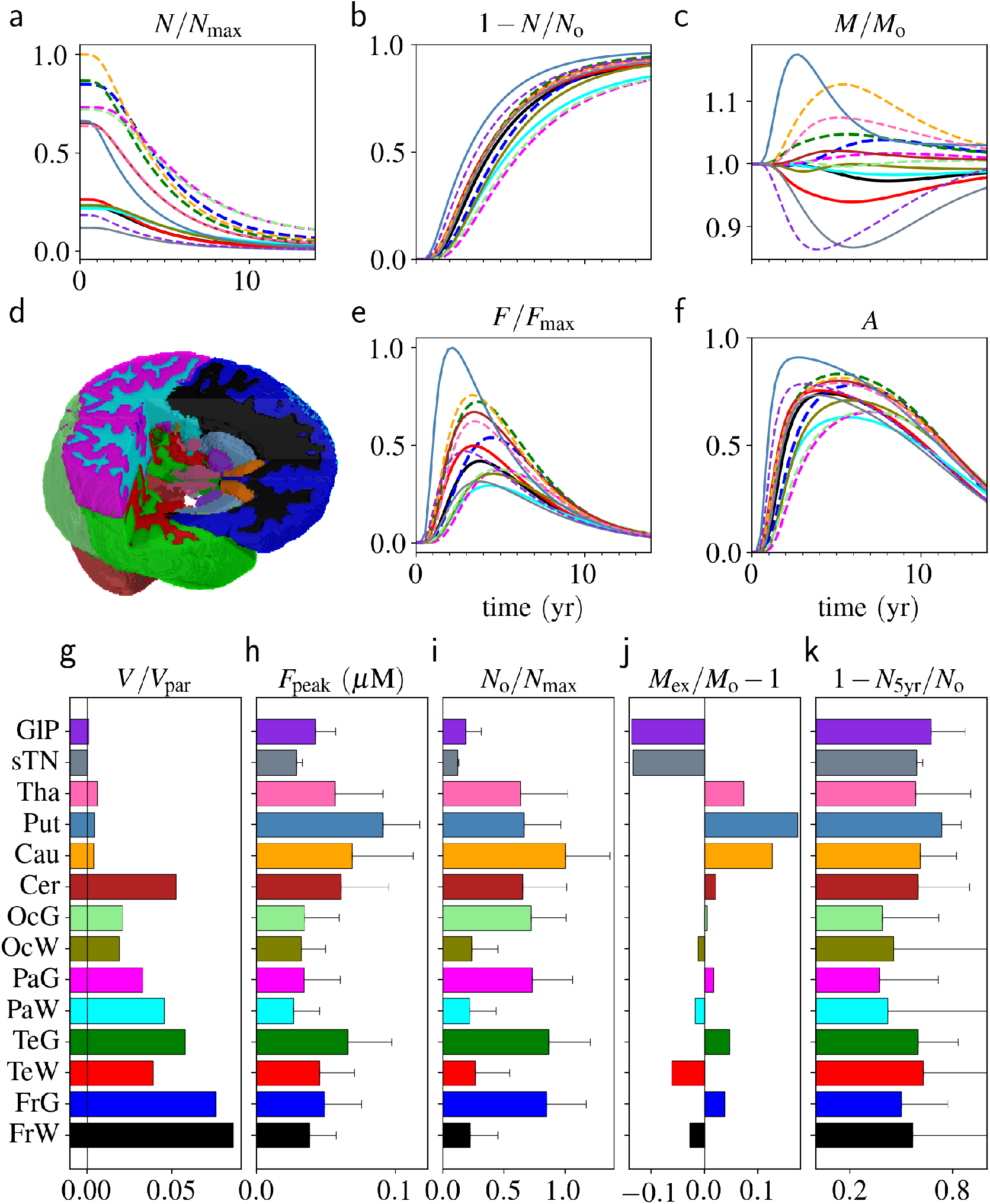
Time evolution of the system variables for different volumes of interest (VOI). Time evolutions are depicted for (a) the neural density *N*(*t*) normalized by the maximum neural density *N*_max_ of the whole brain, (b) atrophy (1-*N/N*_o_ where *N*_o_ = *N*(*t* = 0)), (c) microglial density *M*(*t*) normalized by its initial value (*M*_o_ = *M*(*t* = 0)), (e) amyloid plaque density *F* normalized by the maximum plaque density *F*_max_ of the whole brain, and (f) astrogliosis *A*. (d) A 3D view of different VOIs with the same color-code corresponding to those of all the other panels. (g) The volume fraction of the VOIs in the whole parenchyma. (h) The peak amyloid burden. (i) Original neural density at the initial time (*t* = 0). (j) The peak (extremum) number density of microglia at each VOI. (k) The atrophy *t* = 5 yr. The selected VOIs are abbreviated as the following: FrW: frontal WM, FrG: frontal GM, TeW: temporal WM, TeG: temporal GM, PaW: parietal WM, PaG: parietal GM, OcW: occipital WM, OcG: occipital GM, Cer: cerebellum, Cau: caudate, Put: putamen, Tha: thalamus, sTN: subthalamic nucleus, GlP: globus pallidus.

Studying the time evolution of different system variables in the whole brain is complicated because of its complex structure (see SM1). To deal with this issue, the brain is usually divided into several parts, each of which is called a volume of interest (VOI). The time evolution of the system variables for the most important VOIs provides a clear picture of what happens during the progression of AD (see Fig. 4). A diverse range of A*β* fibrillization, neural death, and brain immune response can be seen among the VOIs. An interesting phenomenon is depicted in Fig. 4-c, demonstrating that microglia concentrate at the more-stressed regions with high A*β* production. This leads to almost uniform atrophy in the brain, as seen in Fig. 4-b. Moreover, the astrogliosis *A* (Fig. 4-f) has a different trend than the microglial activity (Fig. 4-c).

### The movement of microglia and soluble A*β* disrupts the correlation between the neural density and atrophy

If one ignores the spatial derivatives in Eq. 1 and 5, the neural density and atrophy correlate with each other (see supplementary materials). However, the movement of microglia in the brain parenchyma and the propagation of soluble A*β* bring more complexity to the system, leading to the discorrelation between the variables. Accordingly, it is induced from Fig. 4-h, i, and k that the neural density is not always linearly proportional to the maximum amyloid plaque burden or neural atrophy. For instance, although the neural density is almost equal in all the cortex (Frontal GM (FrG), Temporal GM (TeG), Parietal GM (PaG), and Occipital GM (OcG)), they have very different amyloid burden and atrophy. The geometry of the brain adds to this by constraining the movement of microglia and sA*β*. In this regard, the location of CSF and ML plays a crucial role in avoiding or causing amyloidosis and dementia.

The loss of correlation between the system variables could be directly seen by calculating the correlation of z-scores (Fig. 5). Each panel in Fig. 5 shows the correlation map between a pair of system variables at a single time (for the time evolution of correlation, see SM2). From the inset diagrams, it is deduced that the only trivial correlation is between *M* and *F* (Fig. 5-a). Other cross-correlation diagrams denote a completely nonlinear relation between the variables.

**Figure 5:**
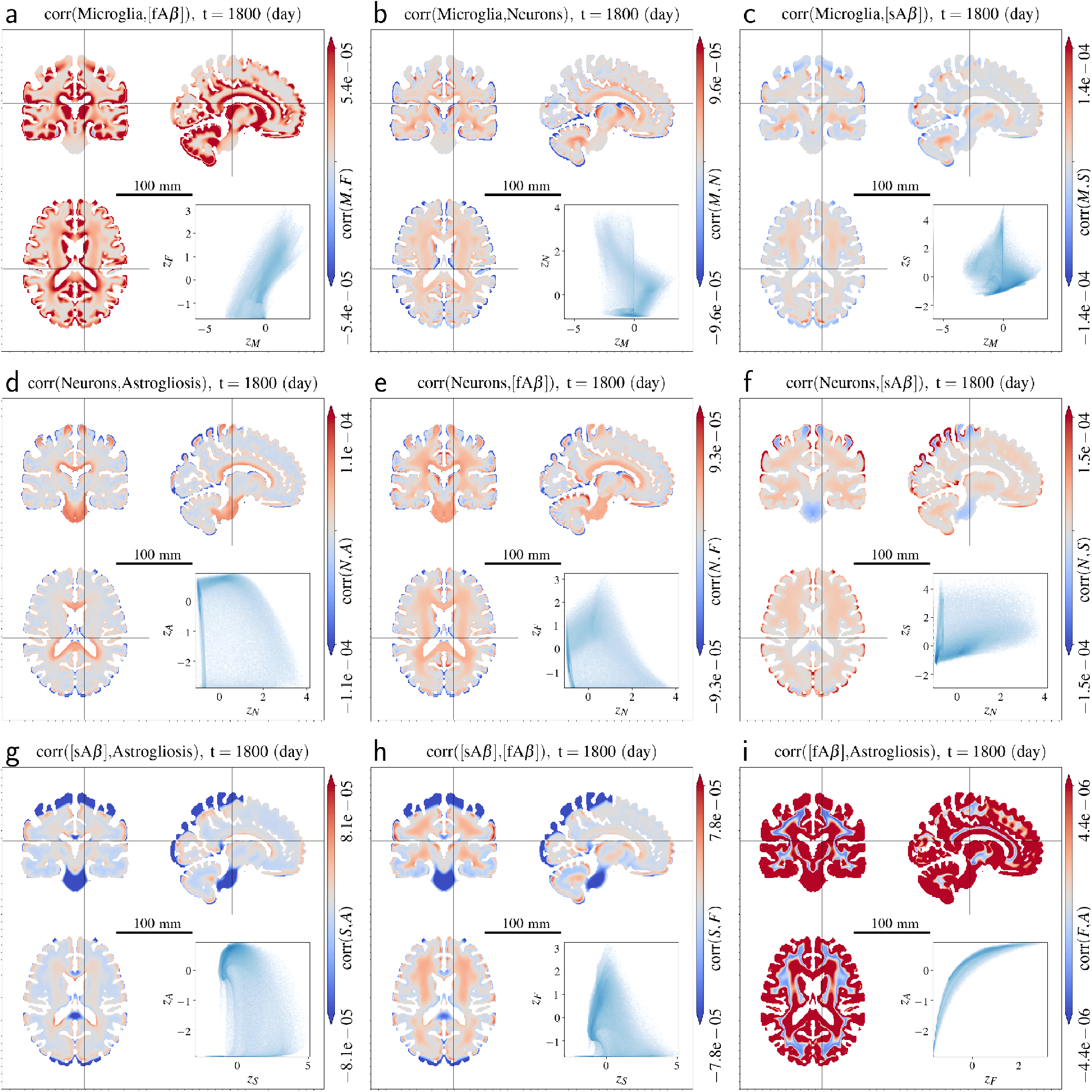
Correlation maps of the z-scores between pairs of system variables at a single time point (*t* = 5 yr). Inset diagrams show the scatter-plots of the variable z-scores in the brain parenchyma. For the time evolution, see SM2. The dependencies between the variables are not simple linear correlations as illustrated by the inset scatter plots. The luminance of the scatter plot is logarithmically normalized.

### ACH is partially inconsistent with the experiments

Comparison of the model predictions with the real AD measurements show consistency in the brain cortex, but not in the central brain (Fig. 6). Since the experimental results are usually taken after death, they do not depict the time evolution of amyloid plaques, neither they show the maximum amount of the plaques. As the model predicts, amyloid plaques get cleared during the course of the disease while neurons die. That means, the maximum amount of amyloid plaques occurs during the progression of the disease rather than at final state. Fig. 6 depicts the state of the amyloid burden at *t* ≈ 10 yr for different brain lobes and compares them with the experimental autopsy results from the work of Arnold et al. [3] in 1991 (indicated by subscript “a”) and McLean et al. [47] in 1999 (indicated by subscript “m”). The predicted results qualitatively agree with the experiments for the brain cortex. However, the predicted amyloid plaques in putamen, thalamus, and cerebellum do not agree with the experiments. This suggests that amyloid plaques are not the only culprits of AD.

**Figure 6:**
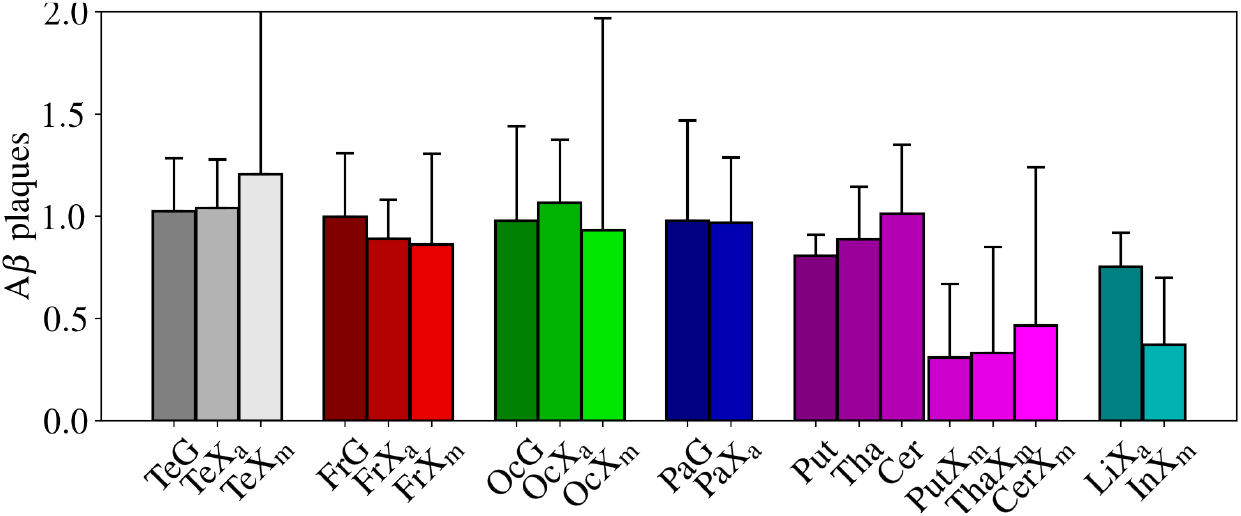
Comparison to experimental results. In silico, the amyloid plaque burden is measured at time *t ≈* 10 yr for each lobe. The experimental results are autopsies (courtesy of Refs. [3, 47]). The simulation and experimental values are normalized based on the average over temporal, frontal, and occipital cortices values. The following abbreviations are used for experimental results; TeX: temporal, FrX: frontal, PaX: parietal, OcX: occipital, LiX: limbic, InX: insular, PutX: putamen, ThaX: thalamus, CerX: cerebellum. subscripts “a” and “m” correspond to the results of Arnold et al. [3] in 1991 and McLean et al. [47] in 1999 respectively.

### Individual differences in Brain Reserve Capacity

In this work, we tried to capture the most probable scenario of AD. However, as many factors in the system are entangled, the variances in the brain properties, from its size and geometry to its immune system strength and neural activity, can substantially alter the time period of the disease as well as the way different parts of the brain respond to the pathology. Here in particular, we focus on the concept of BRC.

For larger original neural density *N*_o_, the time until the neural density reaches a critical threshold increases, even though the amyloid production also increases with the neural density. This statement results directly from the model. Assuming that all the secreted A*β* from neurons transforms to the fibrillar form, the neural death rate ought to be proportional to the production rate of A*β*, 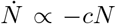. In effect, the neurons would decay exponentially, i.e. *N* = *N*_o_exp(−*t/τ*) with a constant characteristic time *τ*. We can assume that the cognition is affected if the neural density gets smaller than a constant threshold, namely *N*_th_. Comparing two individuals with different *N*_o_, but the same *τ*, the time it takes that the neural density decays to *N*_th_ is longer for the one with greater *N*_o_. With this logic, it is clear why people with higher BRC, due to education or intellectual level [36, 54], or even brain sizes [61], might have lower risk of AD [68].

The critical fibrillization of amyloids with respect to the amyloid production rate [29] and the effect of system parameters on its critical point makes the system very complicated. As shown schematically in Fig. 7, the distribution of the neural density in the whole brain has the most significant effect on amyloid plaque formation and subsequent AD progression. Each individual has a specific distribution of neural density in the brain. In addition, the critical point of fibrillization is also specific to each individual. Hence, three scenarios are possible (Fig. 7): the critical point for fibrillization is outside and above the neural density distribution (case *a*), inside the distribution (case *b*), or outside and below it (case *c*). It is assumed that the system remains healthy below the critical point, but has the risk of AD above it.

**Figure 7:**
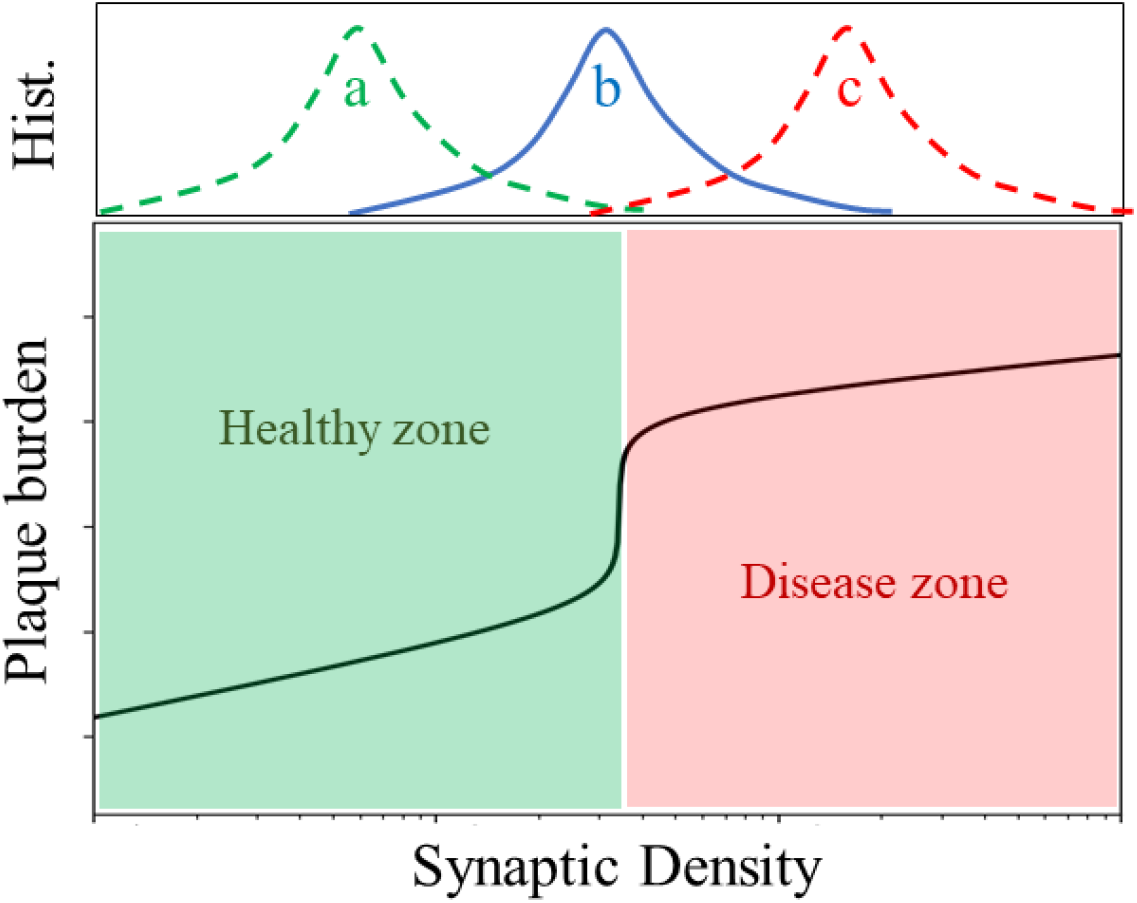
Distribution of neural density in the brain may span (a) only the healthy zone, (b) both healthy and diseased zones, or (c) only the diseased zone. The boundary between the diseased and healthy zone is defined by the system parameters which is specific to each individual.

In this study, case *b* was studied, as the most probable case. Case *b* also results in a diverse atrophy throughout the brain parenchyma. Case *a*, however, should be relevant for healthy individuals since amyloidosis is not prominent. And case *c* is the most extreme case which results in an almost homogeneous atrophy in the brain. Case *b* may also explain why the performance of the individuals in cognitive tests may not correlate with the amount of amyloid plaques and the actual loss of the neural synapses [2]. The amyloid burden and brain atrophy might distribute heterogeneously in the brain according to the distribution of the brain synaptic activity. A heterogeneous atrophy leaves the neuronal network of the brain with multiple pathways for solving cognitive problems. In this regard, cognitive performance is mostly correlated with the distribution of atrophy in the brain rather than with its overall neural density.

### Brain size

The brain size comes into play at the spatial derivative terms in Eq. 1 and 5. These terms could be normalized with respect to the effective radius of the brain, 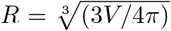 with *V* being the brain volume, by four transformations *X → X/R*^2^ with *X* ∈ *D*_m_, *D*_s_, *ξ*_s_, *ξ*_f_. In comparison, a bigger brain has smaller effective diffusion and chemotaxis, leading to more localized fibrillization and better correlation between the neural density and atrophy. Two distinct cases can be imagined in the context of the brain size. Either the neural density or the total number of neurons remain unchanged with different brain sizes. In the former case, BRC is larger for a bigger brain. In the latter case, neural density is thiner in a bigger brain, leading to lower amyloid production per unit volume. In both cases, a bigger brain results in a later emergence of symptoms, in agreement with clinical evidence [61].

## Conclusion

According to the ACH, amyloid plaque formation is the leading mechanism in AD which triggers a few successive events leading to neural death and dementia [24, 35]. A*β* is secreted out of the neural synapses with neural activity. The higher neural density/activity, the more production of A*β* into the extracellular matrix. Assuming these statements, a system of PDEs has been proposed for modeling the onset and progression of AD.

The model demonstrates that the A*β* production determines whether the produced amyloids accumulate into fibrillar plaques or get discharged out of the brain: If the A*β* production rate in the brain is larger than a critical value (*cN*)_c_, soluble A*β* accumulates into insoluble amyloid plaques, a process taking several years to decades [29]. Assuming that amyloid plaques are both toxic to neurons, and induce neuroinflammation via astrogliosis, their formation leads to the neural death. It implies that another critical amyloid production rate larger than (*cN*)_c_ should exist beyond which neurons start to get toxicated and die. Since the neural density/activity is not homogeneously distributed in the brain, the fibrillization and its subsequent neural death should also be nonhomogeneously distributed.

The amyloid production and their fibrillization in the brain have been addressed in this article. The initial and boundary conditions of the simulations have been translated from MRI data, in order to be as close as possible to reality. The boundary conditions are the tissue structure of the human brain, including the parenchyma and the CSF. The initial conditions of the system stand for the initial neural density which is assumed to be proportional to the intensity of MR images, while WM, GM, and CSF are separated from each other. The simulation results show that some regions of the brain are under higher amyloidosis stress not only due to higher neural density, but also due to other factors, such as their distance to CSF or ML and microglia migration in the parenchyma.

The model shows that the amyloid plaques first form at the regions with highest neural density. Afterwards, microglia migrate toward those regions and clear the formed plaques. At the same time, neurons die because of the plaques and astrogliosis, trigerred by them, and microglial activity. The fibrillization and neural atrophy propagates to other regions of the brain in the later times. The model reveals that soluble A*β* concentration drops significantly both in the brain ISF and CSF as amyloid fibers emerge. The atrophy follows shortly after.

The time evolution of neural death, microglia concentration, fibrillization and astrogliosis have been quantified for the most important volumes of interest (VOIs) in the brain parenchyma. The results indicate that the amyloid plaque formation and clearance peaks at different times for different VOIs. Surprisingly, the peak plaque concentration does not perfectly correlate with the neural density, as one could expect. The plaque formation at the cortex fits well with the experimental findings from autopsies qualitatively. However, the plaque formation at the inner parts of the brain does not match well with the experiments, showing that ACH cannot explain the full process of AD if the amyloidosis is considered as the sole causing factor. This means other factors, such as tau-proteins or neurotoxic inflammation may play an important role in AD. For example, since tau-protein aggregation propagates along the neural fibers, neural death pattern may not be realistic without considering the neural connectome. Nevertheless, such a simplistic view of a complex system, where only the brain geometry and tissue structure are taken into account, predicts the right atrophy and plaque formation in the brain. This implies that the geometry of the brain and its size play crucial roles in the progression of AD.

Finally, the concept of BRC is discussed to be able to explain the diversity of dementia in elderly and the lack of correlation between the amyloidosis and cognitive decline. Individual differences in the brain, including the neural density and activity, the brain geometry, and immune system properties, can lead to a highly diverse pattern of amyloidosis and dementia.

## Acknowledgements

M.H. expresses gratitude to Gang Zhang from the Helmholtz Center for infection Research in Braunschweig, and Vasileios Karakasis at Swiss National Supercomputing Centre for their support of the project. We gratefully acknowledge the HIFIS (Helmholtz Federated IT Services) Software team for support. This project was supported by the Helmholtz-Gemeinschaft, Zukunftsthema “Immunology and Inflammation” (ZT-0027).

## Contributions

M.H. and M.M.H. designed the research project. M.H. derived the model and performed the simulations. J.K., M.Syd., T.M., and M.Scp. supported the simulations. M.H. analyzed the results and wrote the first draft. All authors discussed the results and edited the manuscript. M.M.H. supervised the study.

