## Supplementary information for "Brain geometry matters in Alzheimer’s disease progression: a simulation study"

### Supplementary Materials for "Brain geometry matters in Alzheimer's disease progression: a simulation study"

Masoud Hoore[1] et al.

*

#### Methods

At first, Eq. 1 to 7 in the main text are solved numerically using a mean-field approach in order to understand how the system parameters affect the results. To this end, all the spatial derivatives are omitted and we end up with a set of ODEs.

Afterwards, the full system of PDEs is solved numerically on the brain tissue, taken from the voxel-based structural MRI data of Ref. [16], using a finite-volume method. Eq. 1 to 7 are discretized over the voxels. Fig. S1 schematically shows the setup of such a system. The whole brain structure is read from an MR image with $N_\mathrm{v}^\mathrm{x}$, $N_\mathrm{v}^\mathrm{y}$, and $N_\mathrm{v}^\mathrm{z}$ voxels in sagital ($x$), coronal ($y$), and axial ($z$) directions, respectively. A spatial derivative of a variable for voxel $i$ is translated to the fluxes from voxel $i$ to its adjacent voxels $j(i)$.

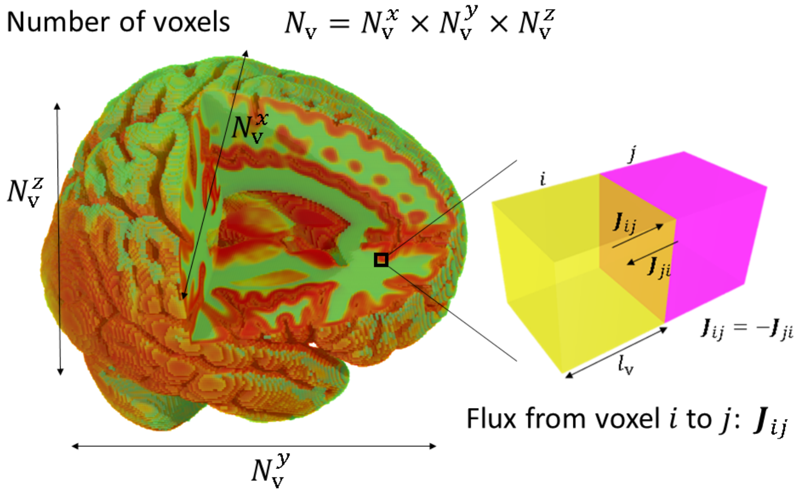


**Figure S1: 3D view of the brain, and a schematic view of the flux calculation for the finite volume method. The color code in the 3D image corresponds to the neural density based on the intensity of the voxel-based structural MRI data from Ref. [16], and the mean neural density in the brain parenchyma estimated from Ref. [5]. The reduced image contains $N_\mathrm{v} = 7109137$ total number of voxels, making a lattice with $N_\mathrm{v}^\mathrm{x} = 181$ voxels in sagital, $N_\mathrm{v}^\mathrm{y} = 217$ voxels in coronal, and $N_\mathrm{v}^\mathrm{z} = 181$ in axial direction. For each pair of voxels, a flux is calculated which represents the spatial derivative of a variable.**

It is noted that there is no accurate data for the neural (synaptic) density in the human brain. Moreover, the neural activity at different regions of the brain is a complicated function of space (i.e. brain regions) and time. In this work, the luminance of the structural MR images "ICBM 152 Nonlinear atlases version 2009" from the McGill Brain Imaging Center (www.bic.mni.mcgill.ca) [16] is taken as a measure of the neural density (see Fig. S6). The luminance has been normalized separately for WM and GM, considering that the neural synaptic density is much smaller in WM than in GM. The neural densities are normalized in the end in order to match the mean neural density $\langle N \rangle = 1.3^7$ mL$^{-1}$.

An open-source software, Braimmu (https://github.com/mhoore/braimmu), has been developed for running the simulations of the equation-based model. Braimmu is written based on C++, using Massage Passing Interface (MPI) and CUDA for parallel computing on CPUs and GPUs. It uses finite-volume method approach for the specific research on the brain immune system. The simulation box is discretized by a cubic lattice with unit size $l_{\mathrm{v}}$. Tab. S1 describes the variables and parameters of the model, with their estimated value due to the provided references.

**Table S1: Model parameters and their estimated values. It is noted that for all the analysis in this work, the parameters are taken from this table unless specified otherwise.**

| Symbol | Description | Estimated value | Ref. |
|---|---|---|---|
| $S$ | concentration of A$\beta$ oligomers in soluble form ([sA$\beta$]) | variable, $\mu$M | |
| $F$ | concentration of A$\beta$ oligomers in fiber form ([fA$\beta$]) | variable, $\mu$M | |
| $M$ | number density of microglia | $\langle M\rangle = 1.1\times10^{7}\pm 11\%\ \mathrm{mL}^{-1}$ | Adapted from [41, 5, 28, 70] |
| $N$ | number density of neurons | $\langle N\rangle = 1.3\times10^{7}\pm 5\%\ \mathrm{mL}^{-1}$ | [5] |
| $A$ | astrogliosis | $0\le A\le 1$ | |
| $c$ | mean rate of sA$\beta$ secretion per neuron | $2\times10^{-11}\ \mu\mathrm{M}\cdot\mathrm{mL}\cdot\mathrm{day}^{-1}$ | Assumed |
| $D_{\mathrm{m}}$ | diffusivity of microglia | $2\times10^{4}\ \mu\mathrm{m}^2\cdot\mathrm{day}^{-1}$ | Rough estimate |
| $\chi_{\mathrm{s}}$ | chemotaxis sensitivity toward sA$\beta$ | $2.2\times10^{4}\ \mu\mathrm{m}^2\cdot\mathrm{day}^{-1}\cdot\mu\mathrm{M}^{-1}$ | Assumed |
| $\chi_{\mathrm{f}}$ | chemotaxis sensitivity toward fA$\beta$ | $2.2\times10^{5}\ \mu\mathrm{m}^2\cdot\mathrm{day}^{-1}\cdot\mu\mathrm{M}^{-1}$ | Assumed |
| $d_{\mathrm{n}}^{\mathrm{a}}$ | neural death rate due to astrogliosis | $1\times10^{-3}\ \mathrm{day}^{-1}$ | Assumed w.r.t [20] |
| $d_{\mathrm{n}}^{\mathrm{f}}$ | neural death rate due to fibrillization | 0; since $A$ is related to $F$, this assumption will not change the behavior of the system significantly. | |
| $H_{\mathrm{a}}$ | Michaelis-Menten constant for astrogliosis | $1\times10^{5}\ \mu\mathrm{M}\cdot\mathrm{mL}^{-1}$ | Assumed |
| $k_{\mathrm{a}}$ | astrogliosis rate | $1\times10^{-2}\ \mathrm{day}^{-1}$ | Assumed |
| $k_{\mathrm{p}}$ | rate of A$\beta$ fiber growth | $2.6\times10^{1}\ \mu\mathrm{M}^{-1}\cdot\mathrm{day}^{-1}$ | Adapted from [13], see [29] |
| $k_{\mathrm{n}}$ | rate of A$\beta$ fiber nucleation | $1.7\times10^{-1}\ \mu\mathrm{M}^{-1}\cdot\mathrm{day}^{-1}$ | Adapted from [13], see [29] |
| $d_{\mathrm{s}}$ | rate of sA$\beta$ clearance by microglia | $5.3\times10^{-9}\ \mathrm{day}^{-1}\cdot\mathrm{mL}$ | [29], [45] and [71, 40], see [29] |
| $e_{\mathrm{s}}$ | rate of sA$\beta$ efflux through the CSF | $1.9\ \mathrm{day}^{-1}$ | [6, 46], see [29] |
| $d_{\mathrm{f}}$ | clearance of fA$\beta$ by microglia | $\sim d_{\mathrm{s}}/30$ | [73], see [29] |

#### Mean-field analysis.

Assuming the mean-field, such a system of equations has only one steady-state solution, which is the trivial solution with $S=F=A=N=0$. The goal is to find out how long it takes for the system to reach its all-dead steady-state. Starting from the initial state

$$\begin{cases} S(t=0)=0 \\ F(t=0)=0 \\ A(t=0)=0 \\ N(t=0)=N_{\mathrm{o}} \end{cases},$$

two scenarios are expected from the system. Note that Eq. 5 drops out in the mean-field approach as $M\equiv$ const. Moreover, in the mean-field approach, CSF/ML, WM, and GM are considered the same, thus $\lambda_{\mathrm{s}}=d_{\mathrm{s}}M+e_{\mathrm{s}}$.

As studied recently for the simple case of fibrillization [29], the concentration of fA$\beta$ is critically affected by the production rate $cN$. Considering that the fibrillization occurs long before the onset of the disease, the level of dementia at the symptomatic stage of AD should be affected similarly by $cN$. Accordingly, the two expected scenarios for the current model depend on $cN$.

On one hand, if the initial A$\beta$ production $cN_{\mathrm{o}}$ is smaller than a critical value, $F$ will not increase much so that it cannot trigger astrogliosis $A$ to increase. Therefore, $A\approx FM/H_{\mathrm{a}}\ll 1$ and the neurons die at a very slow pace. $S$ decays similarly and results in slower production of $F$. The system in this state stays quite stable with a small pace of neural loss. On the other hand, when the production term $cN$ is larger than the critical value, $F$ grows fast to a significant amount and triggers $A$ to fastly grow to unity ($A\to 1$). Consequently, neurons die with a fast pace due to both $F$ and $A$, which leads to less production of $S$ and $F$. Formation of $F$ is blocked in addition because $A\approx 1$ so that $F$ decays, although it is too late for the system to revive because $N$ decays faster than $F$.

As depicted in Fig. S2, the original number of neurons determines how fast the dementia should occur, considering that $c=$ const. Neurons do not decay as long as $cN_{\mathrm{o}}$ is smaller than a critical value $cN_{\mathrm{o}}^{\mathrm{c}}$. When $cN_{\mathrm{o}}>cN_{\mathrm{o}}^{\mathrm{c}}$, the decay starts to happen and gets stronger as $cN_{\mathrm{o}}$ increases.

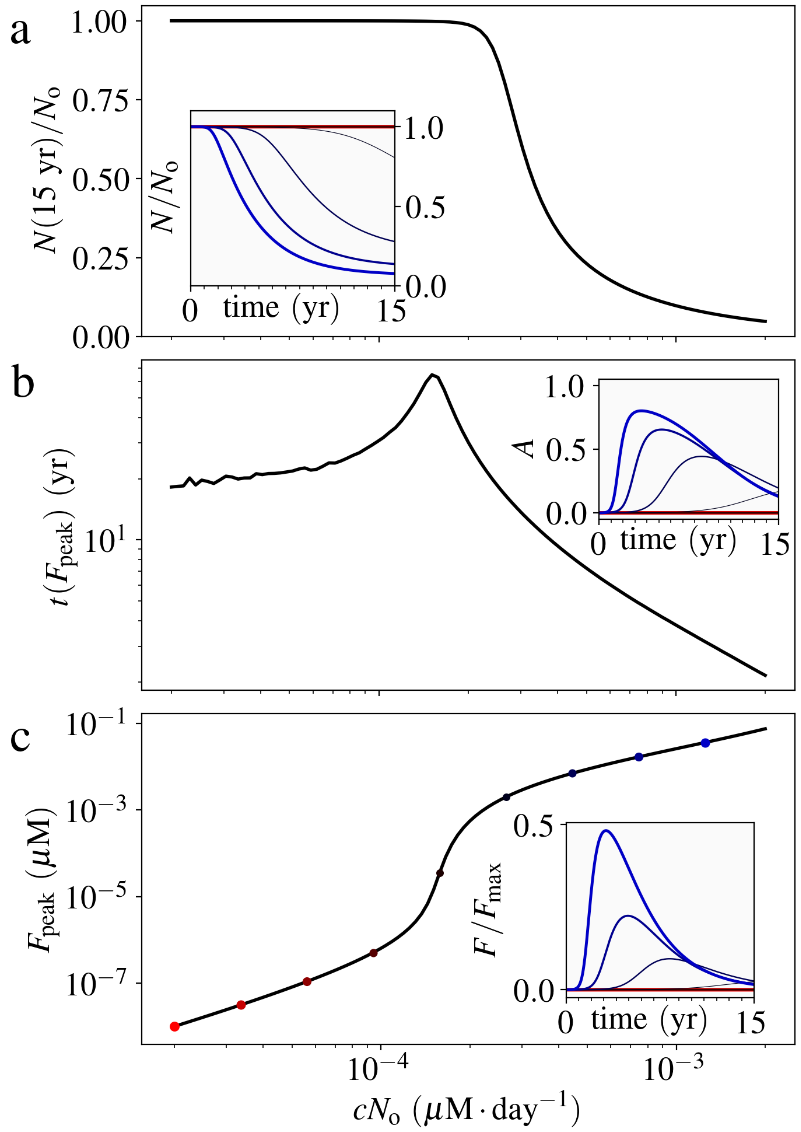


**Figure S2: Effect of the amyloid production on the AD model, provided that the production rate per neuron $c$ is constant in time ($c = \text{const}$). (a) The ratio between the neural density after 15 years and the original value $N(15\ \text{yr})/N_\text{o}$. Neural density $N$ decays if $cN_\text{o}$ surpasses a critical value $cN_\text{o}^\text{c}$. The larger $cN_\text{o}$ is, the faster it decays. (b) The time of reaching the peak of the fibrillized amyloids $F_\text{peak}$, $t(F_\text{peak})$, and (c) the peak fibrillized value $F_\text{peak}$. $cN_\text{o}^\text{c}$ stands for $cN_\text{o}$ at which $t(F_\text{peak})$ peaks or equivalently $F_\text{peak}$ changes abruptly. The inset diagrams of panels a to c show the time evolution of $N/N_\text{o}$, $A$, and $F$, respectively. The initial time $t = 0$ should be assumed as when the homeostatic state of the system is altered. The color codes of the inset diagrams correspond to the markers in panel c with the same color. The parameter values are taken from Tab. S1.**

Similar to the recent studies on the fibrillization process before the onset of AD [29], the system is divided into two states, one with soluble A$\beta$ domination and the other with fibrillar A$\beta$ domination. Astrogliosis $A$ shows a similar pattern as $F$ (see the inset of Fig. S2-b). $N_\text{o}$ and $c$ have the same impact on the system because they both contribute in the production rate similarly. It means there should be a critical value for $c$, too, at which the behavior of the system changes, provided that $c$ remains constant in time. Variations in $c$ at different stages of the disease would lead to more complex results which is out of scope of this work.

As assumed, neural death is driven by two mechanisms: fibrillization and astrogliosis. From Fig. S2, it is deduced that $F$ and $A$ grow together. They are either minuscule, or large and dominant, depending on whether $cN_\text{o}$ is smaller or greater than $cN_\text{o}^\text{c}$. Consequently, the neural death has a similar dependency on $d_\text{n}^\text{f}$ and $d_\text{n}^\text{a}$. Illustrated in Fig. S3, it is implied that $d_\text{n}^\text{a}$ defines how fast neural death takes place. Since the neural density is proportional to the production rate of amyloids, and the production rate is the driving force for the whole system, the decay of neurons affects almost all other variables in the system.

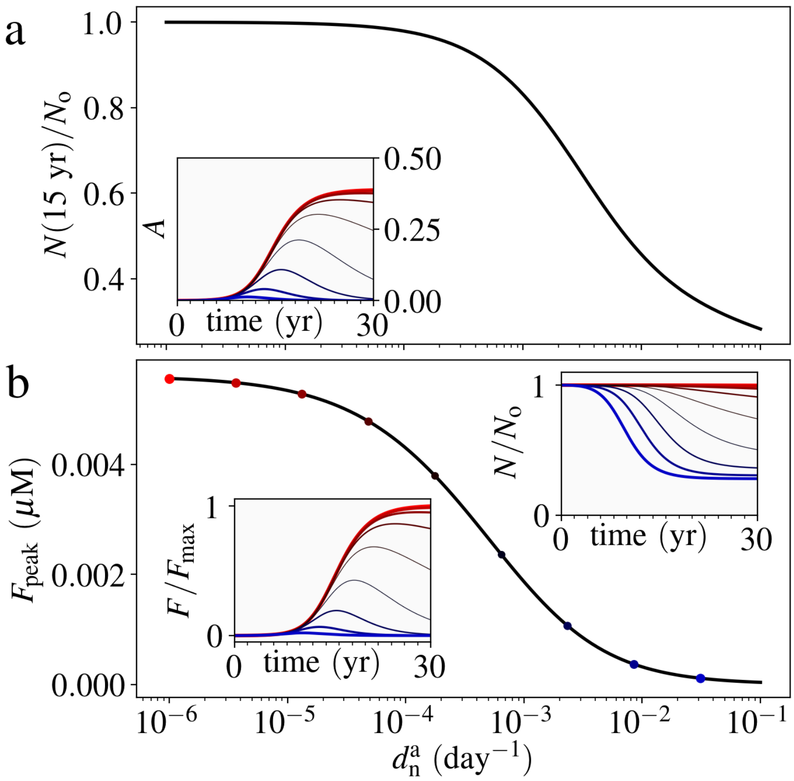


**Figure S3: Effect of the neural death rate due to astrogliosis, $d_n^a$, on the AD model. (a) $N(15\ \mathrm{yr})/N_o$ and (b) $F_{\mathrm{peak}}$ based on $d_n^a$. The inset diagrams show the time evolutions of $N/N_o$, $A$, and $F$ for the corresponding color-coded markers in panel b.**

Fig. S3 shows a similar behavior as Fig. S2. The difference is in the intensities of different responses. Supposing a two-fold state for the system, a healthy and a demented state, the system changes its state with respect to the system parameters. Two ratios are essential in defining such a phase-change behavior in the system, the ratio between the polymerization and nucleation rate $k_p/k_n$, and the ratio between the microglial clearance rate of soluble and fibrillar amyloids $d_s/d_f$ [29]. The physics of the amyloid fibrillization process in the brain parenchyma makes the both ratios very large. Our estimates show that the former ratio is in order of $10^2$ and the latter is in order of $10^1$. Regulating these ratios as an intervention strategy is not a trivial task because it depends on the physical properties of the environment. With this, such a phase-change behavior of the system is universal based on all the other parameters (see also Fig. S4 and Fig. S5).

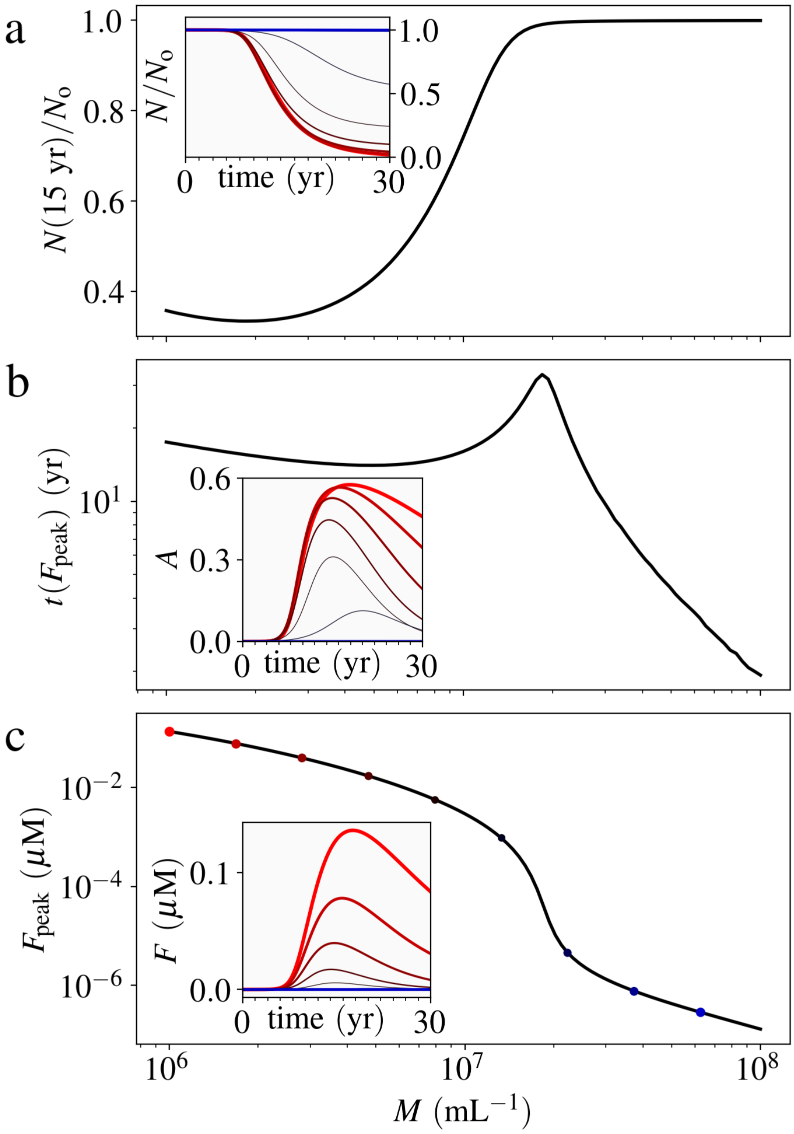


**Figure S4: Effect of the microglial number density $M$ on the AD model. (a) $N(15\ \mathrm{yr})/N_o$, (b) $t(F_{\max})$, and (c) $F_{\max}$ with respect to $M$. Similar to other parameters, there is a critical value for $M$ which separates the diseased and healthy cases. The inset diagrams of panels a to c show the time evolution of $N/N_o$, $A$, and $F$, respectively. The color codes of the inset diagrams correspond to the markers in panel c with the same color.**

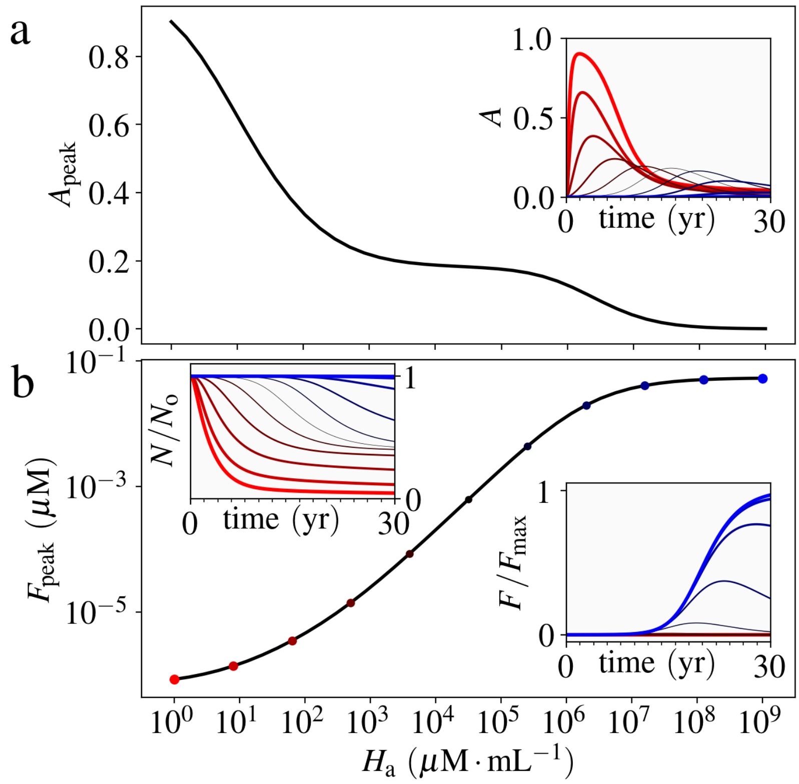


**Figure S5: Effect of the Michaelis constant, $H_{\mathrm{a}}$ for astrogliosis, on the AD model. (a) Maximum astrogliosis $A_{\max}$ and (b) $F_{\max}$ with respect to $H_{\mathrm{a}}$. The inset diagrams show the time evolution of $A$, $N/N_{\mathrm{o}}$, and $F$. The color codes of the inset diagrams correspond to the markers in panel b with the same color.**

#### Other supplementary figures

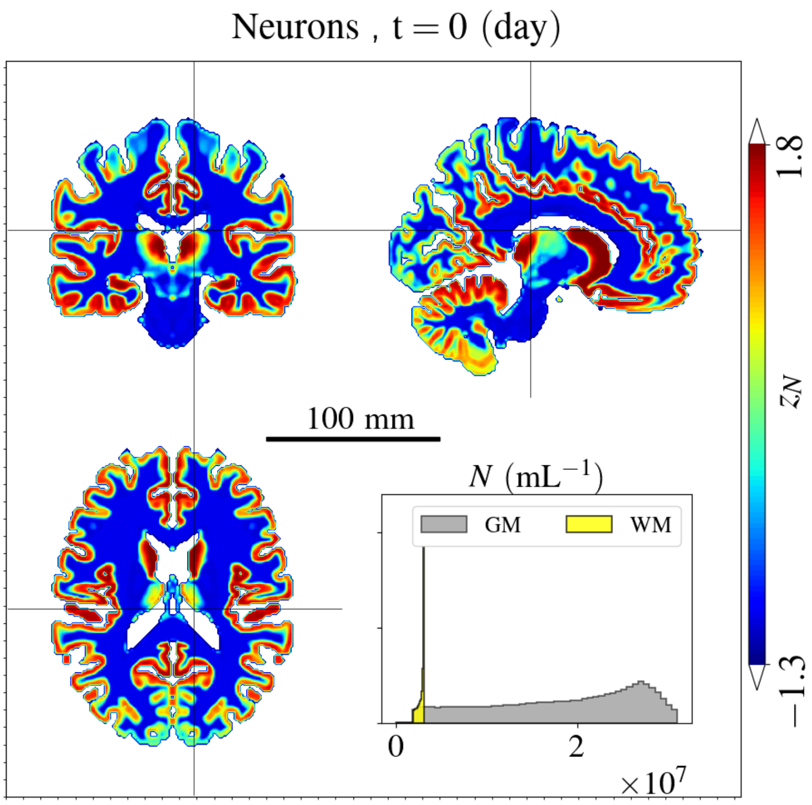


**Figure S6: The normalized neural density (in z-scores) as derived from the luminance of the structural MR images "ICBM 152 Nonlinear atlases version 2009" from the McGill Brain Imaging Center (www.bic.mni.mcgill.ca) [16]. The inset shows the histogram of the neural density.**

#### Supplementary movies

1. SM1: Time evolution of the simulation variables, compare with Fig. 2 and Fig. 3 .
2. SM2: Time evolution of the correlation maps, compare with Fig. 5 .
